# Neonatal gut-microbiome-derived 12,13 DiHOME impedes tolerance and promotes childhood atopy and asthma

**DOI:** 10.1101/311704

**Authors:** S.R. Levan, K.A. Stamnes, D.L. Lin, K.E. Fujimura, D.R. Ownby, E.M. Zoratti, H.A. Boushey, C.C. Johnson, S.V. Lynch

## Abstract

Neonates at risk of childhood atopy and asthma are characterized by gut microbiome perturbation and fecal enrichment of 12,13 DiHOME(1); however, the underlying mechanism and source of this metabolite remain poorly understood. Here we show that 12,13 DiHOME treatment of human dendritic cells altered peroxisome proliferator-activated receptor γ regulated gene expression and decreased immune tolerance. In mice, 12,13 DiHOME treatment prior to airway challenge exacerbated pulmonary inflammation and decreased lung regulatory T cells. Neonatal fecal metagenomic sequencing revealed putative bacterial sources of 12,13 DiHOME. In our cohort, three bacterial genes and their product, 12,13 DiHOME, are associated with increased odds of childhood atopy or asthma, suggesting that early-life gut-microbiome risk factors may shape immune tolerance and identify high-risk neonates years in advance of clinical symptoms.

**One Sentence Summary:** Early-life gut-microbiome risk factors may shape immune tolerance and identify neonates at high-risk of disease.

---

The oxylipin 12,13 DiHOME is significantly enriched in the feces of one-month-old neonates who exhibit a perturbed gut microbiome and are at heightened risk of developing atopy and asthma in childhood (1). In *ex vivo* assays this oxylipin reduces the frequency of regulatory T cells (Tregs) in a dose-dependent manner (1). These observations as well as structural similarities between 12,13 DiHOME and known ligands of peroxisome proliferator-activated receptor γ (PPARγ) (2–5), a nuclear receptor that regulates innate and adaptive immune responses, led us to hypothesize that this oxylipin induces pro-allergic immune dysfunction via PPARγ signaling in dendritic cells (DCs). Accordingly, we treated human DCs with 12,13 DiHOME or vehicle and examined its effects. 12,13 DiHOME treatment decreased DC secretion of IL-10, an anti-inflammatory cytokine that protects against allergic inflammation (6) (Fig. 1a, Fig. S1a). Additionally, co-culture of 12,13 DiHOME-treated DCs with autologous T cells altered the distribution of CD4+ T cells, specifically by decreasing the frequency of Tregs without decreasing cell viability (Fig. 1b-c, Fig. S1b-c).

**Fig. 1.**
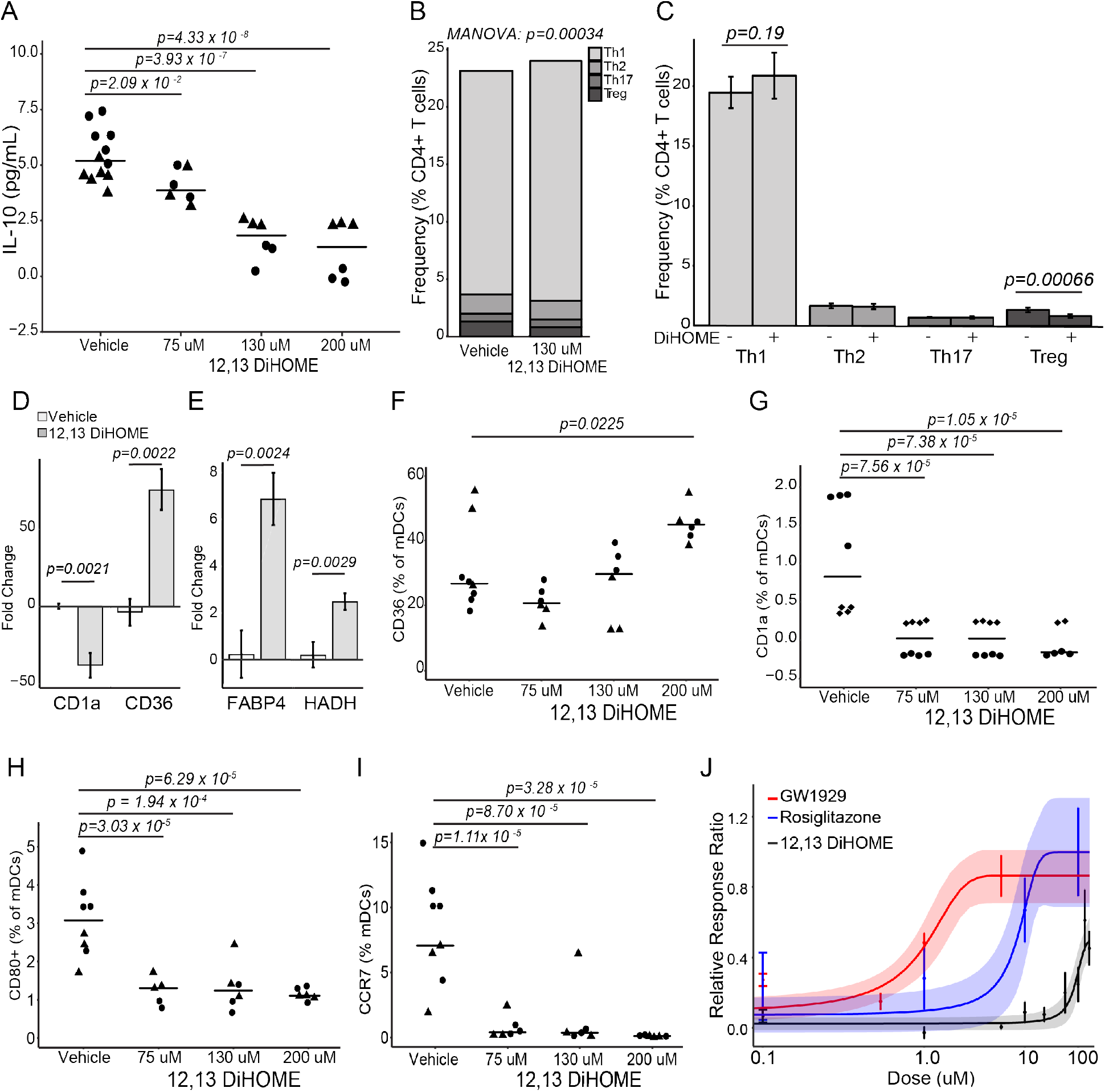
12,13 DiHOME acts via PPARγ on DCs to decrease Tregs. (A) 12,13 DiHOME treatment causes a dose-dependent decrease in IL-10 secretion from human DCs [Linear Mixed Effects (LME); p=0.0209, p=3.93×10^−7^, p=4.33 × 10^−8^ for concentrations of 75, 130, 200 μM, respectively). (B,C) Treatment of DCs with 130 μM 12,13 DiHOME induces a significant shift in the distribution of helper T cells (MANOVA; p=0.00034) and causes a specific decrease in the frequency of Tregs (CD3+CD4+CD25+FoxP3+; LME; p=0.00025). (D,E) Treatment of human DCs with 130 μM 12,13 DiHOME causes changes in gene expression consistent with activation of PPARγ (LME; p=0.0021, p=0.0022, p=0.0024, p=0.0029 for CD1a, CD36, FABP4, and HADH, respectively). (F,G,H,I) 12,13 DiHOME increases the expression of CD36 at high concentrations (LME; p=0.0225 for 200 μM) and decreases the expression of CD1a, CD80, and CCR7 on DCs (CD3-CD19-CD11c+; LME; p<0.001 for all comparisons). Figures 1a-h were performed with n=3-4 treatment replicates using cells isolated from 2 independent donors (biological replicates; ▲, ♦, ●). (J) Raw264.7 cells were transfected with a PPARγ-activated luciferase reporter and treated with 12,13 DiHOME or known PPARγ agonists GW1929 (K_d_ = 1.4 nM) (33) and Rosiglitazone (K_d_ = 40 nM) (34) (n=3). All error bars represent the standard error of the mean (SEM).

To evaluate whether 12,13 DiHOME exerts this effect via PPARγ, we isolated RNA from human DCs treated with 12,13 DiHOME and examined the expression of PPARγ-regulated genes. 12,13 DiHOME treatment mimicked previously characterized effects of PPARγ activation: decreased expression of CD1a, an immune marker involved in lipid presentation (7); increased expression of CD36, a fatty-acid transporter implicated in 12,13 DiHOME-regulation of brown adipocytes (8); and increased expression of FABP4 and HADH, genes involved in fatty acid uptake and metabolism (7) (Fig. 1d-e). To confirm these observations, we used flow cytometry to examine the effects of 12,13 DiHOME on the expression of PPARγ-regulated proteins in human DCs. Again, 12,13 DiHOME treatment mimicked PPARγ activation, increasing expression of CD36, and decreasing expression of CD1a, CD80, and CCR7, immune markers involved in lipid presentation, antigen presentation, and cell trafficking (7), in human DCs (Fig. 1f-i, Fig. S1f-g). To test whether 12,13 DiHOME directly acts on PPARγ, we utilized a modified PPARγ reporter assay (9) and demonstrated that treatment with concentrations of 12,13 DiHOME in excess of 50 μM leads to PPARγ activation (Fig. 1j). While PPARγ activation in DCs is traditionally considered anti-inflammatory (10,11), recent studies suggest that PPARγ may in fact promote allergic inflammation(12). Alternatively, 12,13 DiHOME may act as a weak agonist of PPARγ and compete with endogenous PPARγ ligands found in serum (13).

We next examined whether 12,13 DiHOME exacerbates allergic sensitization *in vivo*, by treating mice with 12,13 DiHOME (30 mg/kg) or vehicle (10% DMSO) via peritoneal injection prior to airway sensitization and challenge with cockroach antigen (CRA) (14). Treated animals exhibit increased peribronchial and perivascular inflammatory infiltrates and serum IgE compared to those treated with vehicle alone (Fig. 2a-d, Fig. S2a-b). 12,13 DiHOME-treated animals also exhibited increases in lung resident T cells, neutrophils, and monocytes, and pulmonary expression of pro-inflammatory innate cytokines, IL1β, IL1α, and TNF, as well as a significant decrease in the frequency of lung Tregs and a trend towards decreased lung resident alveolar macrophages (Fig. 2e-g, Fig. S2g-k). These inflammatory immunologic changes may be explained by the effect of 12,13 DiHOME on PPARγ observed *ex vivo*. Alternatively, 12,13 DiHOME was recently identified as an activator of transient receptor potential vanilloid 1 (TRPVI) (15), loss of which is associated with decreased asthma prevalence in humans and protection against airway inflammation in animal models (16,17). To evaluate whether oxylipins delivered in the peritoneum reach the circulation, we quantified 12,13 DiHOME by liquid-chromatography mass spectrometry (LC-MS) (18) three hours after injection and observed significant enrichment in both the plasma and lungs when compared with vehicle-treated animals (Fig. 2h, Fig. S2l), indicating that 12,13 DiHOME may interact directly with lung tissue to exacerbate allergic airway inflammation.

**Fig. 2.**
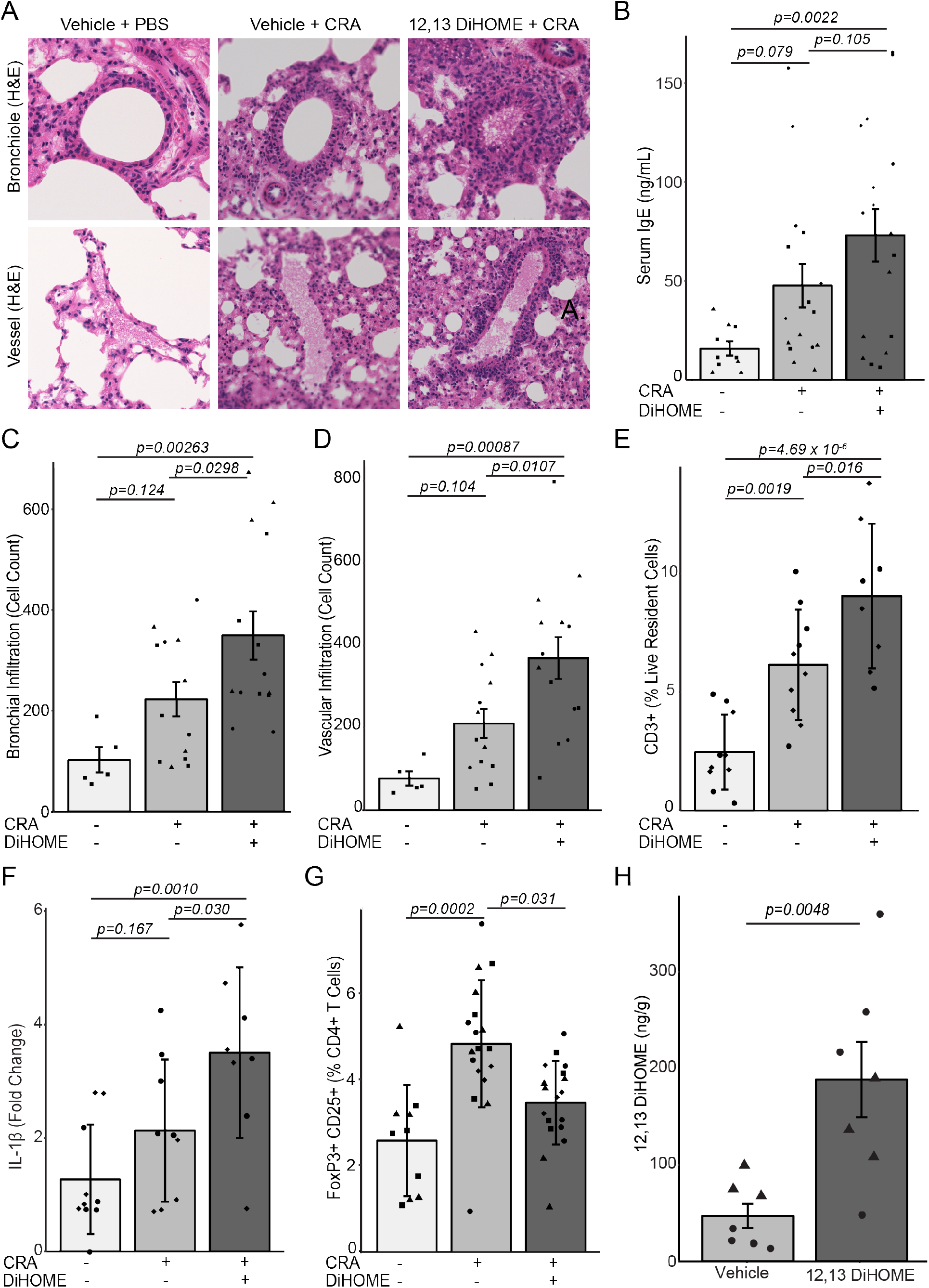
Peritoneal treatment with 12,13 DiHOME exacerbates lung inflammation in CRA-challenged mice. (A) Hematoxylin and eosin stained bronchioles and blood vessels from the lungs of mice treated with vehicle (10% DMSO) and challenged with PBS or CRA or treated with 30 mg kg^−1^ 12,13 DiHOME (solubilized in 10% DMSO) and challenged with CRA. (B) Peritoneal treatment of mice with 12,13 DiHOME (n=17) increases serum IgE compared to CRA-challenged (n=16; LME; p=0.105) and PBS-challenged animals (n=10; LME; p=0.0022). (C,D) 12,13 DiHOME treatment (n=14) increases the number of infiltrating cells surrounding the bronchioles and veins of CRA-challenged mice compared with vehicle treated, PBS challenged [n=5; LME; p=0.00263 (bronchial), p=0.00087 (venous)] or vehicle treated, CRA challenged [n=13; LME; p=0.0298 (bronchial), p=0.0107 (venous)] animals. (E) Peritoneal 12,13 DiHOME treatment increases the frequency of lung resident T cells (CD3+) in CRA challenged mice (n=9) compared with vehicle treated, PBS challenged (n=10; LME; p=4.69×10^−6^) or vehicle treated, CRA challenged (n=10; LME; p=0.016) animals. (F) 12,13 DiHOME treatment (n=8) increases the expression of IL1β compared with vehicle treated, PBS challenged (n=9; LME; p=0.0010) or vehicle treated, CRA challenged (n=9; LME; p=0.030) animals. (G) 12,13 DiHOME treatment of CRA-challenged mice (n=18) decreases Treg frequency compared to vehicle-treated CRA-challenged mice (n=18; LME; p=0.031). (H) Peritoneal treatment with 12,13 DiHOME significantly increases the concentration of 12,13 DiHOME in the lungs (n=6; two-sided student t-test: p=0.0048) at 3 hours post-delivery. Unique symbols (▲, ▀, ♦, ●) represent mice from independent assays. All error bars represent the SEM.

Given the apparent role of this lipid in driving pro-allergic immune dysfunction in mice and humans, we focused on determining the concentration and potential sources of 12,13 DiHOME in neonatal feces. We began by quantifying 12,13 DiHOME and 9,10 DiHOME (its enantiomer and a known agonist of PPARγ (19)) using LC-MS in a subset of one-month-old neonates from the Wayne County Health, Environment, Allergy & Asthma Longitudinal Study (WHEALS; Table 1; total n=41; atopic=7; asthmatic=8; atopic asthmatic=4) (1). These included 26 one-month old stool samples that had been previously selected for untargeted metabolomic profiling based on their representation of the varied gut microbiome composition observed across 130 one-month old subjects (1), and an additional 15 randomly selected one-month old samples that had more than 50 mg of stool and 10 ng of extracted fecal DNA remaining, and had childhood atopy and/or asthma outcomes available. 12,13 DiHOME was present in all neonatal stool, but was detected at significantly higher concentrations in the stool of neonates who subsequently developed atopy and/or asthma (Fig. 3a). In contrast the concentration of 9,10 DiHOME did not significantly differ between the two groups (Fig. S3a).

12,13 DiHOME is a terminal metabolic product of linoleic acid, which is initially converted to 12,13 EpOME either enzymatically via a cytochrome P450 epoxygenase (20) or spontaneously via oxidation; 12,13 EpOME is then converted to 12,13 DiHOME via an epoxide hydrolase (EH); an enzyme encoded by humans, bacteria and fungi (21–23). To identify potential sources of 12,13 DiHOME, the aforementioned 26 neonatal stool samples (1) underwent shotgun metagenomic sequencing. A database of known bacterial (~73,000), fungal (~5,000), and human (~50) EH genes was assembled and used in conjunction with ShortBred (24) to probe the neonatal metagenomic data for sequence reads with EH homology. No fungal or human EH genes were detected; however, approximately 1,400 bacterial EH genes were identified. Bacterial EH genes were significantly more abundant in the stool of neonates who developed atopy and/or asthma (Fig. 3b). The thirty most abundant of which were primarily encoded by *Enterococcus faecalis, Streptococcus, Bifidobacterium bifidum*, and *Lactobacillus* strains (Fig. 3c).

**Fig. 3.**
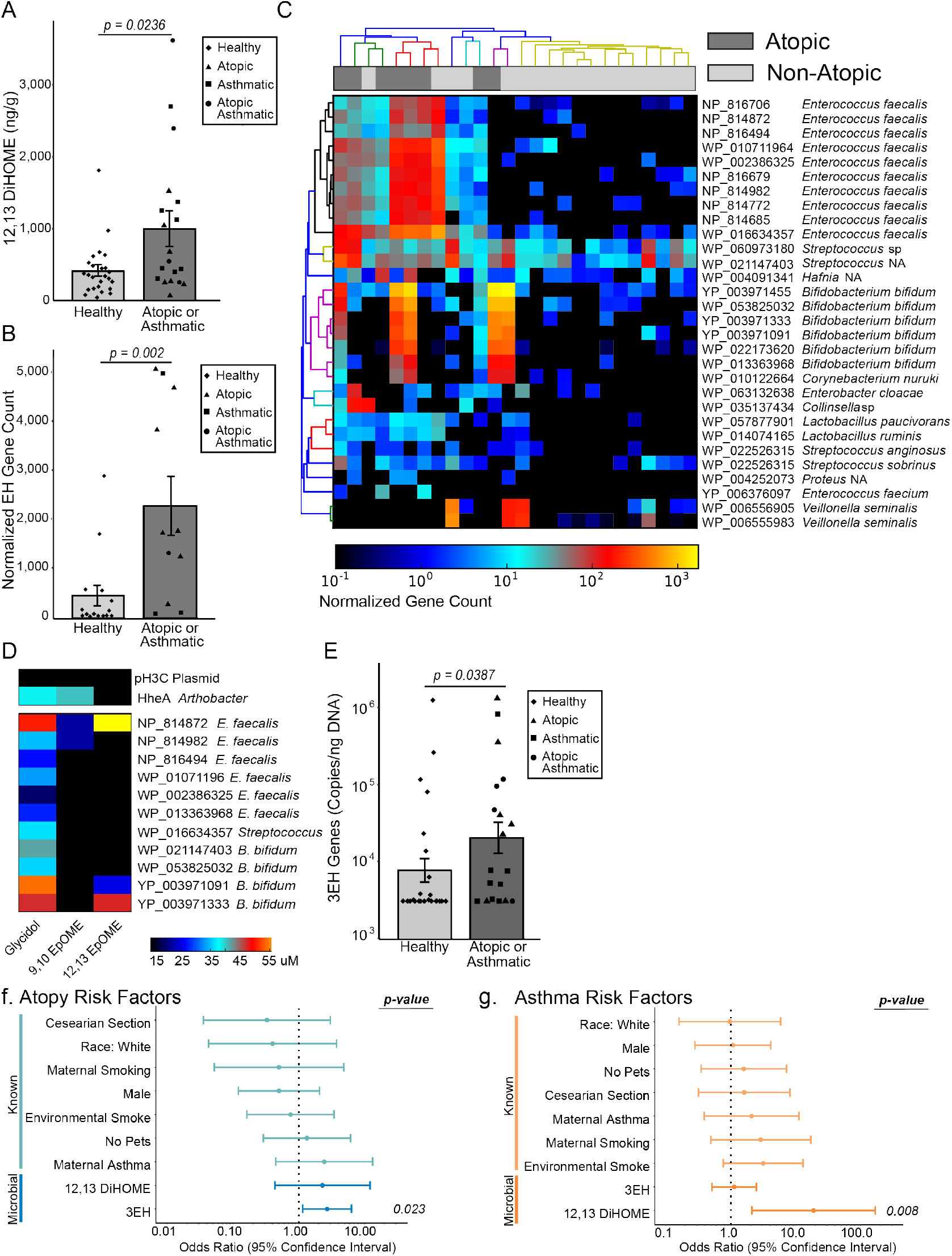
*Enterococcus faecalis* and *Bifidobacterium bifidum* strains in the neonatal gut microbiome encode the capacity to produce 12,13 DiHOME and increase the odds of developing atopy at age two. (A) Neonates who develop childhood atopy and/or asthma exhibit increased fecal concentrations of 12,13 DiHOME (n=41; two-sided Wilcox test; p=0.0236). (B) Neonates who develop atopy at age two have more bacterial EH genes in their stool (n=26; twosided Wilcox test; p=0.002). (C) The 30 most abundant EH genes identified in stool are enriched in neonates who develop atopy. (D) Three of the candidate EH genes, *E. faecalis* NP_814872, *B. bifidum* YP_003971091, and *B. bifidum* YP_003971333, convert 12,13 EpOME to its conjugate diol, 12,13 DiHOME. (E) 3EH copy number mimics the significant increase in overall fecal EH genes observed in neonates who go on to develop atopy and/or asthma (n=41; two-sided Wilcox test; p=0.0387). (F) Increased 3EH copy number in neonatal stool significantly increased the relative odds of developing atopy at age two [n=41; Univariate Logistical Regression; Odds ratio (OR)=2.70; 95% confidence interval (CI)= 1.15-6.36; p=0.023]. (G) Increasing 12,13 DiHOME concentrations in neonatal stool significantly increased the relative odds of developing asthma at age four [n=41; Univariate Logistical Regression; OR=18.8; CI=2.12-166; p=0.008].

**Table 1.** Comparison of sample characteristics for atopy at age two and asthma at age four. P values were calculated using a Fisher Exact test for all comparisons except age at the time of stool collection, which was calculated using a two-sided student T-test.

|  | Comparisons for atopy at age 2 |  |  | Comparisons for asthma at age 4 |  |  |
| --- | --- | --- | --- | --- | --- | --- |
|  | Atopic<br>(n=11) | Non-atopic<br>(n=30) | P value | Asthmatic<br>(n=12) | Non-asthmatic<br>(n=29) | P value |
| Race |  |  |  |  |  |  |
| White, no. (%) | 1 (2.4) | 6 (15) | 0.651 | 2 (4.9) | 5 (12) | 1.000 |
| Black, no. (%) | 9 (22) | 23 (56) | 1.000 | 9 (22) | 23 (56) | 1.000 |
| Male, no. (%) | 4 (9.8) | 16 (39) | 0.484 | 6 (15) | 14 (34) | 1.000 |
| Age of Stool,<br>median (SD) | 35 (4.24) | 34.5<br>(21.5) | 0.280 | 35 (14.3) | 35 (20.5) | 0.830 |
| Maternal Asthma,<br>no. (%) | 3 (7.3) | 4 (9.8) | 0.361 | 3 (7.3) | 4 (9.8) | 0.398 |
| Cesarean section,<br>no. (%) | 1 (2.4) | 7 (17) | 0.412 | 3 (7.3) | 5 (12) | 0.672 |
| Exclusively<br>Breastfed, no. (%) | 1 (2.4) | 3 (7.3) | 1.000 | 0 (0) | 4 (9.8) | 0.302 |
| Maternal Smoking,<br>no. (%) | 1 (2.4) | 5 (12) | 1.000 | 3 (7.3) | 3 (7.3) | 0.334 |
| Environmental<br>Smoke Exposure,<br>no. (%) | 3 (7.3) | 10 (24) | 1.000 | 6 (15) | 7 (17) | 0.146 |
| No Pet Exposure,<br>no. (%) | 8 (20) | 20 (49) | 1.000 | 9 (22) | 19 (46) | 0.719 |

To evaluate whether these putative EH genes are capable of hydrolyzing epoxides and producing 12,13 DiHOME, a cell-based assay was developed. A subset (n=11) of the most frequently detected putative bacterial EH genes, with ≥75% of homologous EH marker regions identified in the metagenomic data, were selected (Table S1). These genes were synthesized and cloned into *Escherichia coli* for expression. EH activity was measured using modifications of a previously described colorimetric assay (25). All 11 genes are capable of hydrolyzing glycidol, a generic epoxide, to its conjugate diol, glycerol, and three can hydrolyze 9,10 EpOME to 9,10 DiHOME. However, only NP_814872 (*E. faecalis*), YP_003971091 (*B. bifidum*), and YP_003971333 (*B. bifidum*) can convert 12,13 EpOME into 12,13 DiHOME (Fig. 3d), indicating that while EH activity is common, the specific capacity to produce 9,10 or 12,13 DiHOME may be bacterial strain specific.

To test whether the three 12,13 DiHOME-producing bacterial EH genes (3EH) were sufficient to distinguish neonates who developed atopy and/or asthma in childhood from those who did not, we developed qPCR assays to quantify the 3EH copy number in neonatal stool. We measured fecal 3EH copy number in our subset of the WHEALS cohort and observed a significant increase in the stool of neonates who subsequently developed atopy and/or asthma in childhood (Fig. 3e). Armed with these data we evaluated whether fecal 12,13 DiHOME and 3EH copy number were associated with increased odds of childhood atopy or asthma. Using our subset of the WHEALS cohort (n=41, Table 1) and univariate logistical regression, we examined the odds of developing atopy at age two years or asthma at age four years based on neonatal fecal 12,13 DiHOME concentration, fecal 3EH copy number, or known early-life risk factors selected *a priori* based on previously published literature. These included male gender; race; maternal asthma; lack of pets, maternal smoking, and environmental smoke exposure pre-delivery; delivery by Cesarean section; and formula feeding at one month (26–32).

Univariate modeling indicated a significant increase in the odds of atopy at age two years with increased fecal 3EH, and a significant increase in the odds of asthma at age four years with an increased fecal concentration of 12,13 DiHOME at one month (Figure 3f,g, Table S6,7). Multivariate modeling was used to test the remaining risk factors for potential confounding. There was no strong evidence for confounding in the relationship between 3EH copy number and the development of atopy at age two (all changes in odds ratio < 20%); however, when we examined the relationship between fecal 12,13 DiHOME concentration and the development of asthma at age four, race was identified as a potential confounder. We adjusted for race in our model and found that 12,13 DiHOME concentration remained significantly associated with asthma development at age four (Table S7). Clearly these neonatal gut microbiome risk factors require validation in additional larger cohorts. Nonetheless they offer potential for early life identification of those at risk for atopy and asthma years in advance of clinical symptoms, when prevention is most likely to be effective.

## Acknowledgments

We would like to acknowledge the WHEALS study participants, Dr. Anthony Iavarone and the mass spectrometry facility and genomic sequencing laboratory at QB3 Berkeley (http://qb3.berkeley.edu/), and generous plasmid donations by Drs. Oren Rosenberg, Bert Vogelstein, and Bruce Spiegelman. Nicolas Lukacs for critical assessment of the manuscript.

## Author contributions

SRL designed the study, performed research, developed the manuscript; KAS, performed research and contributed to manuscript development; DLL and KEF, performed research; DRO, EMZ and CCJ provided samples and data from the WHEALS cohort and contributed to manuscript development; HAB contributed to manuscript development; SVL, designed the study, developed the manuscript

## Data and materials availability

Metagenomic data generated in this study is available in the EMBLI repository Accession # PRJEB24006 (https://www.ebi.ac.uk/). Additional datasets and materials are available from the corresponding author upon request. Data sets and R scripts used for statistical analysis and figures are available on GitHub (https://github.com/srlevan/).

## Supplementary Files

Materials and Methods Supplementary Text Figs. S1 to S5 Tables S1 to S7

## References and Notes

1. K. E. Fujimura et al., Neonatal gut microbiota associates with childhood multisensitized atopy and T cell differentiation. Nat. Med. 22, 1187–1191 (2016).

2. V. N. Vangaveti et al., Hydroxyoctadecadienoic Acids Regulate Apoptosis in Human THP-1 Cells in a PPARγ-Dependent Manner. Lipids. 49, 1181–1192 (2014).

3. M. X. Byndloss et al., Microbiota-activated PPAR-γ signaling inhibits dysbiotic Enterobacteriaceae expansion. Science. 357, 570–575 (2017).

4. A. Khare, K. Chakraborty, M. Raundhal, P. Ray, A. Ray, Cutting Edge: Dual Function of PPARγ in CD11c+ Cells Ensures Immune Tolerance in the Airways. J. Immunol. 195, 431–435 (2015).

5. W. Wahli, L. Michalik, PPARs at the crossroads of lipid signaling and inflammation. Trends in Endocrinology & Metabolism. 23, 351–363 (2012).

6. S. S. Iyer, G. Cheng, Role of interleukin 10 transcriptional regulation in inflammation and autoimmune disease. Crit. Rev. Immunol. 32, 23–63 (2012).

7. I. Szatmari et al., PPAR regulates the function of human dendritic cells primarily by altering lipid metabolism. Blood. 110, 3271–3280 (2007).

8. M. D. Lynes et al., The cold-induced lipokine 12,13-diHOME promotes fatty acid transport into brown adipose tissue. Nat. Med. 37, 1685 (2017).

9. J. Choo et al., A Novel Peroxisome Proliferator-activated Receptor (PPAR)γ Agonist 2-Hydroxyethyl 5-chloro-4,5-didehydrojasmonate Exerts Anti-Inflammatory Effects in Colitis. J. Biol. Chem. 290, 25609–25619 (2015).

10. G. Woerly et al., Peroxisome proliferator-activated receptors alpha and gamma down-regulate allergic inflammation and eosinophil activation. J Exp Med. 198, 411–421 (2003).

11. H. Hammad et al., Activation of peroxisome proliferator-activated receptor-gamma in dendritic cells inhibits the development of eosinophilic airway inflammation in a mouse model of asthma. The American Journal of Pathology. 164, 263–271 (2004).

12. S. P. Nobs et al., PPARγ in dendritic cells and T cells drives pathogenic type-2 effector responses in lung inflammation. J Exp Med. 8, jem.20162069 (2017).

13. I. Szatmari et al., PPAR regulates the function of human dendritic cells primarily by altering lipid metabolism. Blood. 110, 3271–3280 (2007).

14. K. E. Fujimura et al., House dust exposure mediates gut microbiome Lactobacillus enrichment and airway immune defense against allergens and virus infection. Proc. Natl. Acad. Sci. U.S.A. 111, 805–810 (2014).

15. D. Green et al., Central activation of TRPV1 and TRPA1 by novel endogenous agonists contributes to mechanical allodynia and thermal hyperalgesia after burn injury. Mol Pain. 12, 174480691666172 (2016).

16. Q. Wang et al., [TRPV1 UTR-3 polymorphism and susceptibility of childhood asthma of the Han Nationality in Beijing]. Wei Sheng Yan Jiu. 38, 516–521 (2009).

17. K. Baker et al., Role of the ion channel, transient receptor potential cation channel subfamily V member 1 (TRPV1), in allergic asthma. Respiratory Research. 17, 143 (2016).

18. S. Gouveia-Figueira et al., Mass spectrometry profiling of oxylipins, endocannabinoids, and N-acylethanolamines in human lung lavage fluids reveals responsiveness of prostaglandin E2 and associated lipid metabolites to biodiesel exhaust exposure. Anal Bioanal Chem. 409, 2967–2980 (2017).

19. B. Lecka-Czernik et al., Divergent effects of selective peroxisome proliferator-activated receptor-gamma 2 ligands on adipocyte versus osteoblast differentiation. Endocrinology. 143, 2376–2384 (2002).

20. J. Ha, M. Dobretsov, R. C. Kurten, D. F. Grant, J. R. Stimers, Effect of linoleic acid metabolites on Na(+)/K(+) pump current in N20.1 oligodendrocytes: role of membrane fluidity. Toxicol. Appl. Pharmacol. 182, 76–83 (2002).

21. C. Morisseau, Role of epoxide hydrolases in lipid metabolism. Biochimie. 95, 91–95 (2013).

22. B. K. Biswal et al., The molecular structure of epoxide hydrolase B from Mycobacterium tuberculosis and its complex with a urea-based inhibitor. J. Mol. Biol. 381, 897–912 (2008).

23. M. Decker, M. Arand, A. Cronin, Mammalian epoxide hydrolases in xenobiotic metabolism and signalling. Arch. Toxicol. 83, 297–318 (2009).

24. J. Kaminski et al., High-Specificity Targeted Functional Profiling in Microbial Communities with ShortBRED. PLoS Comput Biol. 11, e1004557 (2015).

25. L. Tang et al., A high-throughput adrenaline test for the exploration of the catalytic potential of halohydrin dehalogenases in epoxide ring-opening reactions. Biotechnology and Applied Biochemistry. 62, 451–457 (2015).

26. G. Wegienka et al., Combined effects of prenatal medication use and delivery type are associated with eczema at age 2 years. Clin. Exp. Allergy. 45, 660–668 (2015).

27. G. Wegienka et al., Subgroup differences in the associations between dog exposure during the first year of life and early life allergic outcomes. Clin. Exp. Allergy. 47, 97–105 (2017).

28. S. Havstad et al., Effect of prenatal indoor pet exposure on the trajectory of total IgE levels in early childhood. J. Allergy Clin. Immunol. 128, 880–885.e4 (2011).

29. M. J. Gallant, A. K. Ellis, What can we learn about predictors of atopy from birth cohorts and cord blood biomarkers? Ann. Allergy Asthma Immunol. 120, 138–144 (2018).

30. H. Burke et al., Prenatal and passive smoke exposure and incidence of asthma and wheeze: systematic review and meta-analysis. Pediatrics. 129, 735–744 (2012).

31. Y. Bao et al., Risk Factors in Preschool Children for Predicting Asthma During the Preschool Age and the Early School Age: a Systematic Review and Meta-Analysis. Curr Allergy Asthma Rep. 17, 85 (2017).

32. C. J. Lodge et al., Perinatal cat and dog exposure and the risk of asthma and allergy in the urban environment: a systematic review of longitudinal studies. Clinical and Developmental Immunology. 2012, 176484–10 (2012).

33. Brad R Henke et al., N-(2-Benzoylphenyl)-l-tyrosine PPARγ Agonists. 1. Discovery of a Novel Series of Potent Antihyperglycemic and Antihyperlipidemic Agents. Journal of Medicinal Chemistry. 41, 5020–5036 (1998).

34. J. M. Lehmann et al., An antidiabetic thiazolidinedione is a high affinity ligand for peroxisome proliferator-activated receptor gamma (PPAR gamma). J. Biol. Chem. 270, 12953–12956 (1995).

